# ABO genetic variation in Neanderthals and Denisovans

**DOI:** 10.1101/2020.07.27.223628

**Authors:** Fernando A. Villanea, Emilia Huerta-Sanchez, P. Keolu Fox

## Abstract

Variation at the ABO locus was one of the earliest sources of data in the study of human population identity and history, and to this day remains widely genotyped due to its importance in blood and tissue transfusions. Here, we look at ABO blood type variants in our archaic relatives: Neanderthals and Denisovans. Our goal is to understand the genetic landscape of the ABO gene in archaic humans, and how it relates to modern human ABO variation. We found two derived Neanderthal variants of the O allele in the Siberian Neanderthals (O1 and O2), one of these variants is shared with an European Neanderthal, who is a heterozygote for this O1 variant and a rare cis-AB variant. The Denisovan individual is heterozygous for two ancestral variant of the O1 allele, similar to variants found widely in modern humans. Perhaps more surprisingly, the O2 allele variant found in Siberian Neanderthals can be found at low frequencies in modern Europeans and Southeast Asians, and the O1 allele variant found in Siberian and European Neanderthal is also found at very low frequency in modern East Asians. Our genetic distance analyses suggest both alleles survive in modern humans due to inbreeding with Neanderthals. We find that the sequence backgrounds of the surviving Neanderthal-like O alleles in modern humans retain a higher sequence divergence than other surviving Neanderthal genome fragments, supporting a view of balancing selection operating in the Neanderthal ABO alleles by retaining highly diverse haplotypes compared to portions of the genome evolving neutrally.

## 2 Introduction

ABO was the first blood group system discovered in humans, its identity coded by the ABO gene. As one of the first genetic markers, variation at the ABO gene has been studied for over 60 years, and yet there are some aspects of its evolution that remain mysterious. In early years, ABO blood types could be identified through chemical assays, so ABO gene variation became important as a source of genetic information before the advent of sequencing technology. For example, Cavalli-Sforza and Mordant used serological-based determination of ABO blood types in the 1960s to determine if ABO allele frequencies accurately recapitulated historic human migration patterns [Cavalli-Sforza et al., 1964]. Advances in sequencing technology allowed for improved resolution of ABO locus variation, including detection of population-specific rare variants. Rare variants are useful as a tool to understand population migration history. Examples include a rare O variant (O1V542) which is an Ancestry Informative Marker in Indigenous American populations [Estrada-Mena et al., 2010, Villanea et al., 2013], as well as myriad other rare variants private to specific human groups [Yip, 2002, Roubinet et al., 2004]. In addition, Fry et al. [2007] speculates that rare variation is also informative about natural selection due to historical interactions with pathogens such as malaria, norovirus, smallpox, and perhaps others, further highlighting the importance of studying rare ABO variation.

In modern humans, common variation at the ABO gene is determined by very few non-synonymous polymorphisms. The ABO gene codes for a glycosyltransferase which modifies a precursor O antigen into the A or B antigens. Only four mutations change the enzymatic function of the glycosyltransferase responsible for the A or B antigens. A single deletion can prevent glycosyltransferase enzymatic function such that O antigens are not catalyzed into A or B. These common A, B, and O alleles maintain higher-than-expected level of haplotype diversity - a classic signature of balancing selection [Ségurel et al., 2012, Seymour et al., 2004, Villanea et al., 2015]. ABO haplotypes have common functional types (i.e., A, B, and O), which are retained in populations by the effects of balancing selection, but at a finer scale, rare variation accumulates through mutation, creating diverging haplotype backgrounds for alleles of near identical functional types (i.e., A1, A2, etc). For archaic humans - Neanderthals and Denisovans - the recent sequencing of high-coverage genomes opens up the possibility of studying what ABO diversity looks like in these extinct species.

We now know that when anatomically modern humans dispersed outside of Africa, they encountered and hybridized with various archaic human groups [Green et al., 2010]. Direct comparisons of archaic and modern human genomes have revealed a complex landscape of admixture between modern humans and both Neanderthals and Denisovans [Browning et al., 2018, Villanea and Schraiber, 2019]. For some genes found in modern humans, negative natural selection has acted on archaic variants removing them from the gene pool [Sankararaman et al., 2016, Petr et al., 2019, Zhang et al., 2020], other archaic variants may have been lost through the effects of genetic drift and population bottlenecks. However, a handful of archaic versions of genes have risen to high frequency in modern humans through positive natural selection [Huerta-Sánchez et al., 2014, Huerta-Sánchez and Casey, 2015, Racimo et al., 2015, 2016]. It is possible then, that archaic ABO variants could have been inherited by modern humans, and become targets of natural selection in modern human populations.

Here, we investigate population-specific rare variation of the ABO gene by combining data from the 1,000 Genomes Project, the Neanderthal genome project [Prü fer et al., 2014], the Chagyrskaya Neanderthal genome project [Mafessoni et al., 2020], and the Denisovan Genome Project [Meyer et al., 2012]. Our naive expectation for ABO archaic alleles is to find common functional types with analog function to the modern human A, B, and O alleles, as well as for mutation to give rise to unique archaic haplotype variants. We find that the Denisovan individual presents functional variants that are similar to modern ABO haplotypes found in African and non-African living individuals, possibly retained in both lineages since before the Denisovan-human split. The Altai, Chagyrskaya, and Vindija individuals all present unique Neanderthal ABO haplotypes. Furthermore, these Neanderthal variants are found in select human non-African populations as a result of human-Neanderthal admixture (Figure 1). Finally, we find that the high sequence divergence observed in the introgressed Neanderthal ABO haplotypes is consistent with the effects of balancing selection.

**Figure 1:**
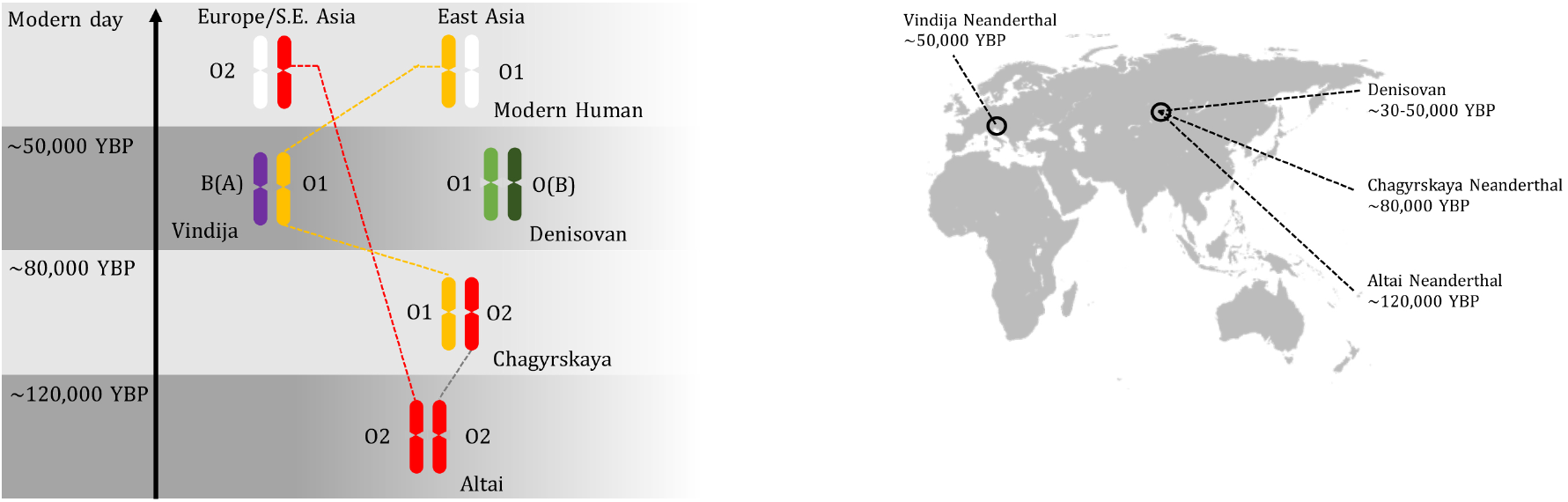
Archaic ABO haplotype sharing through time, and geographic location for three Neanderthal and one Denisovan individuals. The O2 allele found in the earlier Altai Neanderthal is found in modern humans in Europe and Southeast Asia. The Chagyrskaya Neanderthal presents the same O2 allele (but different haplotype background) and a O1 allele shared with the later Vindija Neanderthal, and found in modern humans East Asia. The Denisovan O variants are not directly shared with either Neanderthals or modern humans.

## 3 Results

### 3.1 ABO variation in the 1000 Genomes Project

In modern humans, common ABO allele variation occurs in exons 6 and 7 of the ABO gene, where five common SNPs and two deletions determine the common allele types. We extracted those positions for all archaic haplotypes, as summarized in Table 1. Changes in the first five positions alter the function of the glycosyltransferase responsible for A1 (ABO*A101), A2 (ABO*A201), or B1 (ABO*B101) alleles. The common O1 (ABO*O01) and O1v (ABO*O02) alleles are determined by the last two positions on Table 1, where the last position, a loss-of-function (LOF) deletion, halts enzyme function completely [Yamamoto et al., 1992]. The less common O2 (ABO*O03) allele typically lacks the (LOF) deletion, yet present no glycosyltransferase activity, the result of an mutation producing a complete but non-functional glycosyltransferase.

**Table 1:**
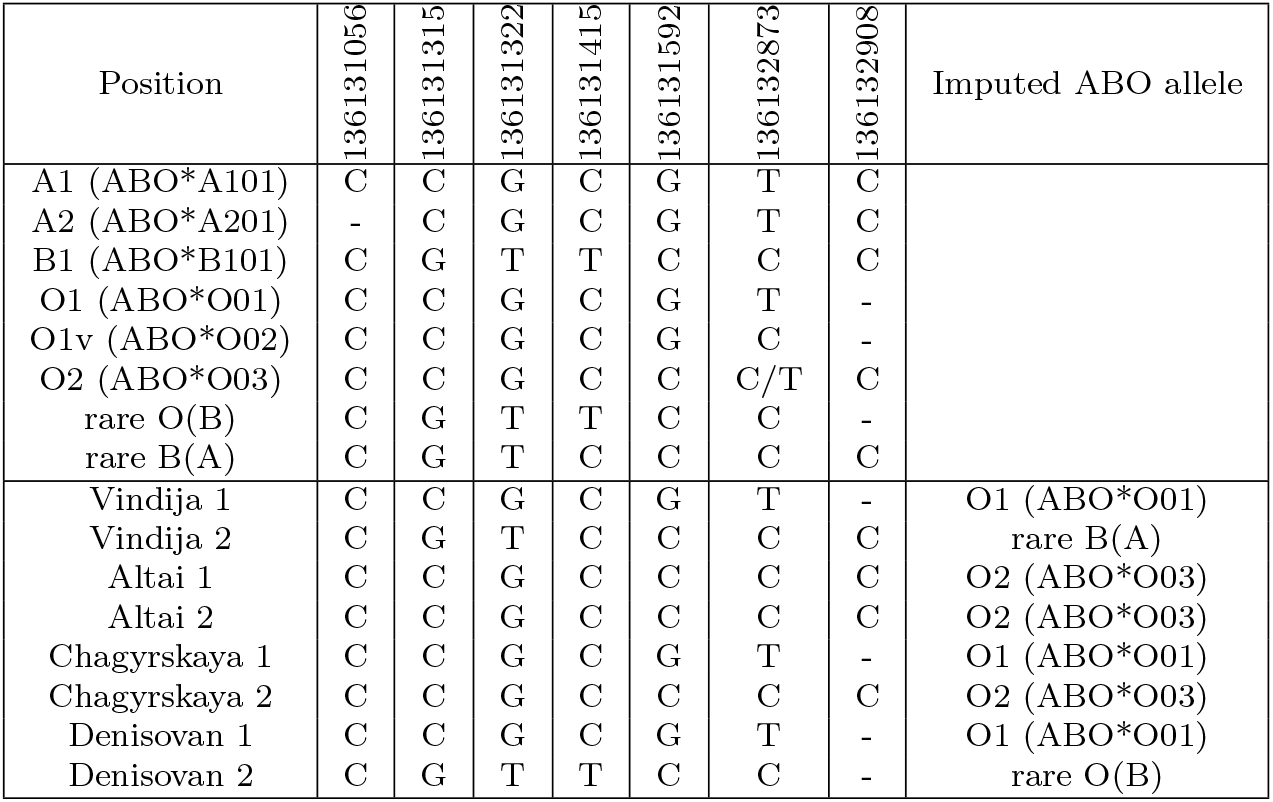
Genotypes for the 6 most common ABO variants and 2 rare variants, including chromosome 9 position in the human genome (Hg19, build37), as well as genotypes for each chromosome in the 4 archaic individuals. Imputed ABO allele types for the four archaic individuals based on these positions are shown on the right-most column.

Then, to understand archaic ABO functional variation at the rare haplotype level, we summarized the ABO variation in 2,504 diploid individuals in the 1,000 Genomes Project panel (the coding sequence of 5,008 chromosomes), identifying 68 variable sites, and resulting in a total of 108 unique haplotypes: 12 unique A1 alleles (703 chromosomes), 8 unique A2 alleles (242 chromosomes), 9 unique B alleles (720 chromosomes), 40 unique O1 alleles (1838 chromosomes), 29 unique O1v alleles (1404 chromosomes), 6 unique O2 alleles (56 chromosomes), 3 rare O (B background) alleles (44 chromosomes), and finally, a single rare cis-AB allele (1 chromosome, Figure 2).

**Figure 2:**
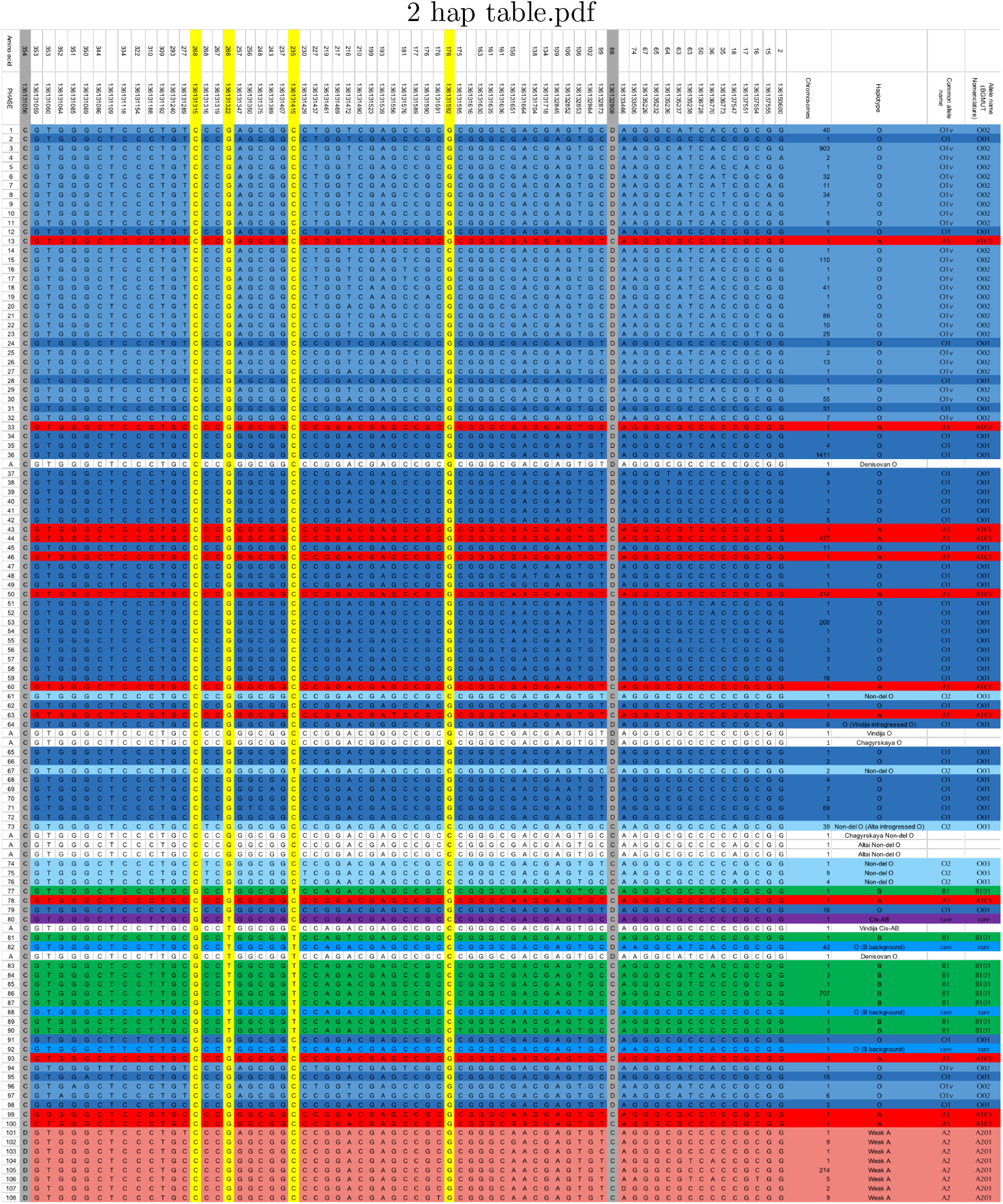
Annotated PHASE output for the 1,000 Genomes Project and archaic genomes. 114 unique haplotypes total. Lower-case bases indicate missing data in the archaic genomes imputed from the modern data.

### 3.2 Archaic human ABO haplotype structure

Using the coding variation at the ABO locus in the 1000 Genomes Project panel as a reference, we identified ABO blood types for four archaic individuals (Table 1, Supp. Figure 1). The single Denisovan individual presents two forms of O alleles (both presenting the typical LOF deletion at position Hg19 9:136132908). The first allele, a typical 01 is identical to 1411 chromosomes in the 1000 Genomes Project, while the second O allele is identical to a rare O (B background) allele found in 42 chromosomes in the 1000 Genomes Project. The Altai Neanderthal individual is homozygous for a rare O2 haplotype [Yip, 2002], different from those found in 39 modern humans chromosomes by a single mutation at position Hg19 9:136131316. The Chagyrskaya Neanderthal individual was also heterozygous for the same O2 haplotype found in Altai, and an O1 allele found in 6 modern human chromosomes, and shared with the Vindija Neanderthal. The Vindija Neanderthal individual is heterozygous for an O1 allele, shared with Chagyrskaya, and a functional allele similar to a rare cis-AB (B(A)) modern human allele found in a single individual in the 1000 Genomes Panel (HG02537), different only at position Hg19 9:136131461. Cis-AB alleles are exceedingly rare recombinant alleles presenting features of both A and B glycosyltransferases in a single chromosome [Chun et al., 2019].

### 3.3 Genetic distance between archaic and human ABO haplotypes

To identify if the archaic ABO haplotypes are related to the modern versions found in the 1000 Genomes Project individuals, we calculated sequence divergence between ABO haplotypes (hg19 coordinates, 9:136130563-136150630) using Haplostrips. Haplostrips is a tool designed to sort and cluster haplotypes based on the distance between the genetic sequences, and ordering them with respect to a reference, in this case an archaic individual. Distances in Haplostrips are Manhattan distances, simply the number of SNPs with different alleles in two sequences. Finally, haplotypes are reordered by decreasing similarity with the archaic reference haplotype. We then visualized these distances grouped by modern human population (Figure 3, Figure 4). Note that admixed populations were excluded from this analysis, the justification can be found in the methods, and Supp. Table 3. Our results indicate that all archaic haplotypes have close equivalents in modern humans. In the case of Denisovans, their O1 and O(B) alleles are close to other O alleles common in modern populations, in particular O alleles found in African individuals. This suggests that the Denisovan O allele variants are ancestrally shared between Denisovans and modern humans. This is not unlikely, considering balancing selection has maintained the same ABO functional alleles for millions of years among primate species [Sé gurel et al., 2012].

**Figure 3:**
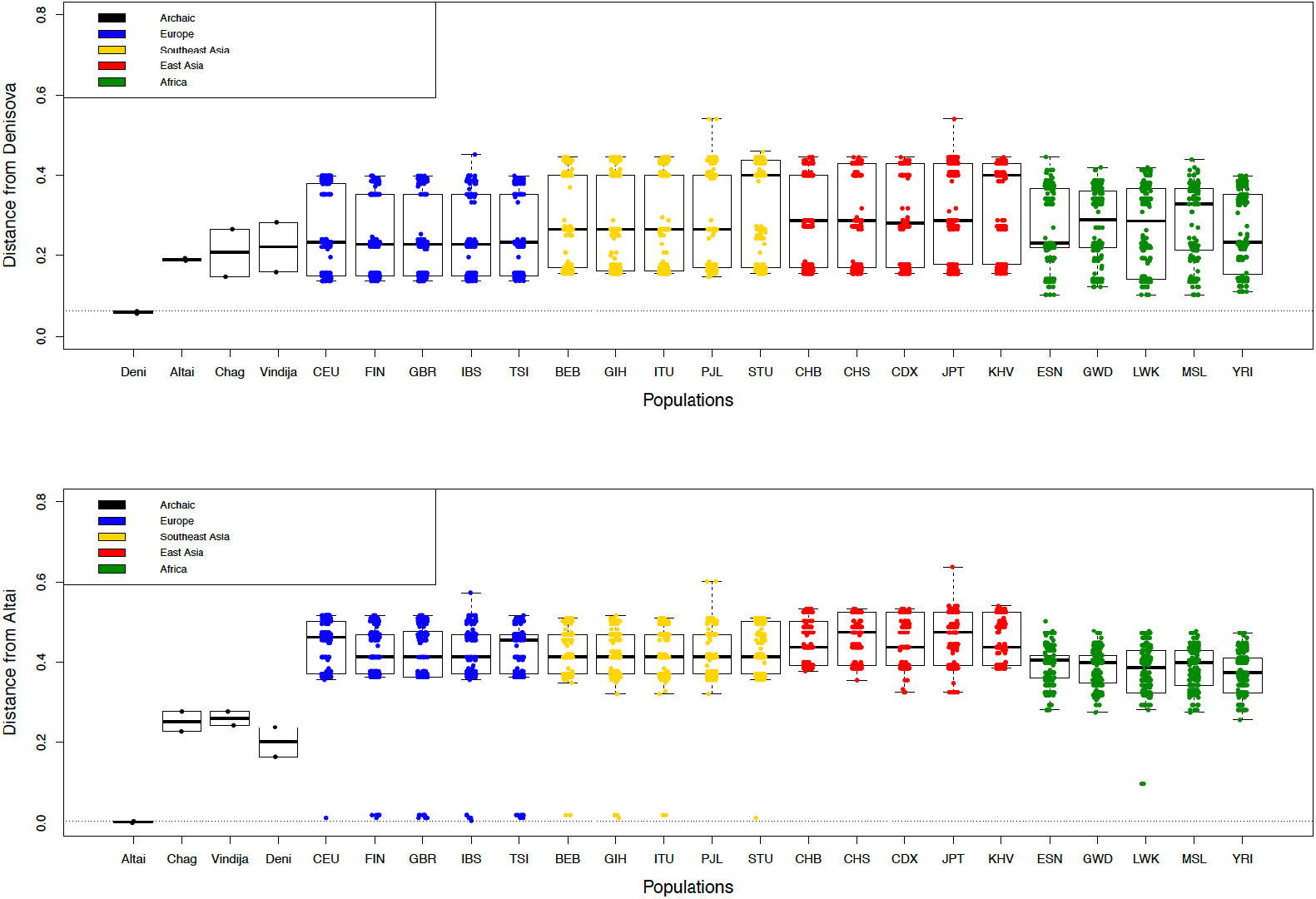
Haplotype distances calculated in Haplostrips for the 1000 Genomes Project populations, polarized relative to the archaic allele: top) Denisova, bottom) Altai Neanderthal. Note that archaic haplotypes are unphased, and heterozygous alleles are randomly assigned between the two chromosomes.

**Figure 4:**
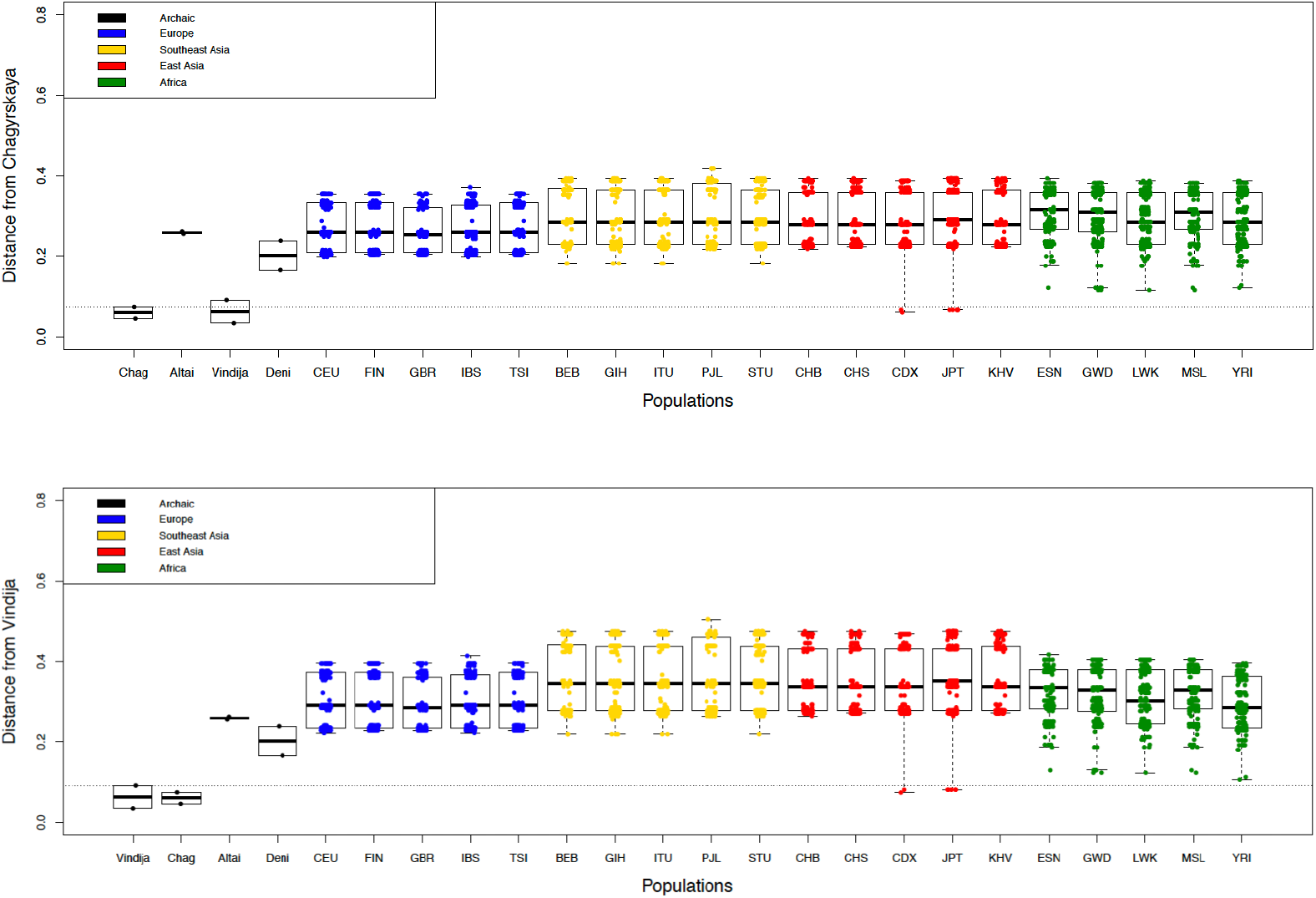
Haplotype distances calculated in Haplostrips for the 1000 Genomes Project populations, polarized relative to the archaic allele: top) Chagyrskaya Neanderthal, bottom) Vindija Neanderthal. Note that archaic haplotypes are unphased, and heterozygous alleles are randomly assigned between the two chromosomes.

Conversely, for Neanderthals, both the Altai/Chagyrskaya O2 and Vindija/Chagyrskaya O1 alleles have almost identical haplotype equivalents in modern humans in non-African populations. The geographic pattern of these alleles is a compelling case for introgression, the result of admixture between Neanderthals and modern humans in Europe, Southeast Asia, and East Asia.

To confirm if the Neanderthal-like ABO alleles found in Europe, Southeast Asia, and East Asia are the result of introgression, we searched in the list of introgressed genome fragments reported in Browning et al. [2018]. We found introgressed genome fragments that overlap with the genome coordinates of the ABO gene in seven populations in the 1000 Genomes Project panel (Supp. Table 1), supporting introgression as the likely cause of shared ABO haplotypes between Neanderthals and modern humans.

### 3.4 Introgressed Neanderthal ABO haplotypes have different affinities compared to all other introgressed fragments genome-wide

We found two different introgressed Neanderthal O haplotypes in modern humans, an Altai/Chagyrskaya-like O2 allele found in modern Europeans and Southeast Asians, and a Vindija/Chagyrskaya-like O1 allele found in modern East Asians. Given this complex geographic pattern, as the Vindija Neanderthal is from Croatia and Altai and Chagyrskaya Neanderthals are from Siberia, we wanted to compare the affinity of ABO introgressed haplotypes in modern humans to the Vindjia (y-axis) and Altai (x-axis) Neanderthal genomes, relative to the affinity of all other introgressed fragments in the genomes of a European (IBS) and East Asian (JPT) population. For this purpose, we generated an affinity contour density plot (Figure 5).

**Figure 5:**
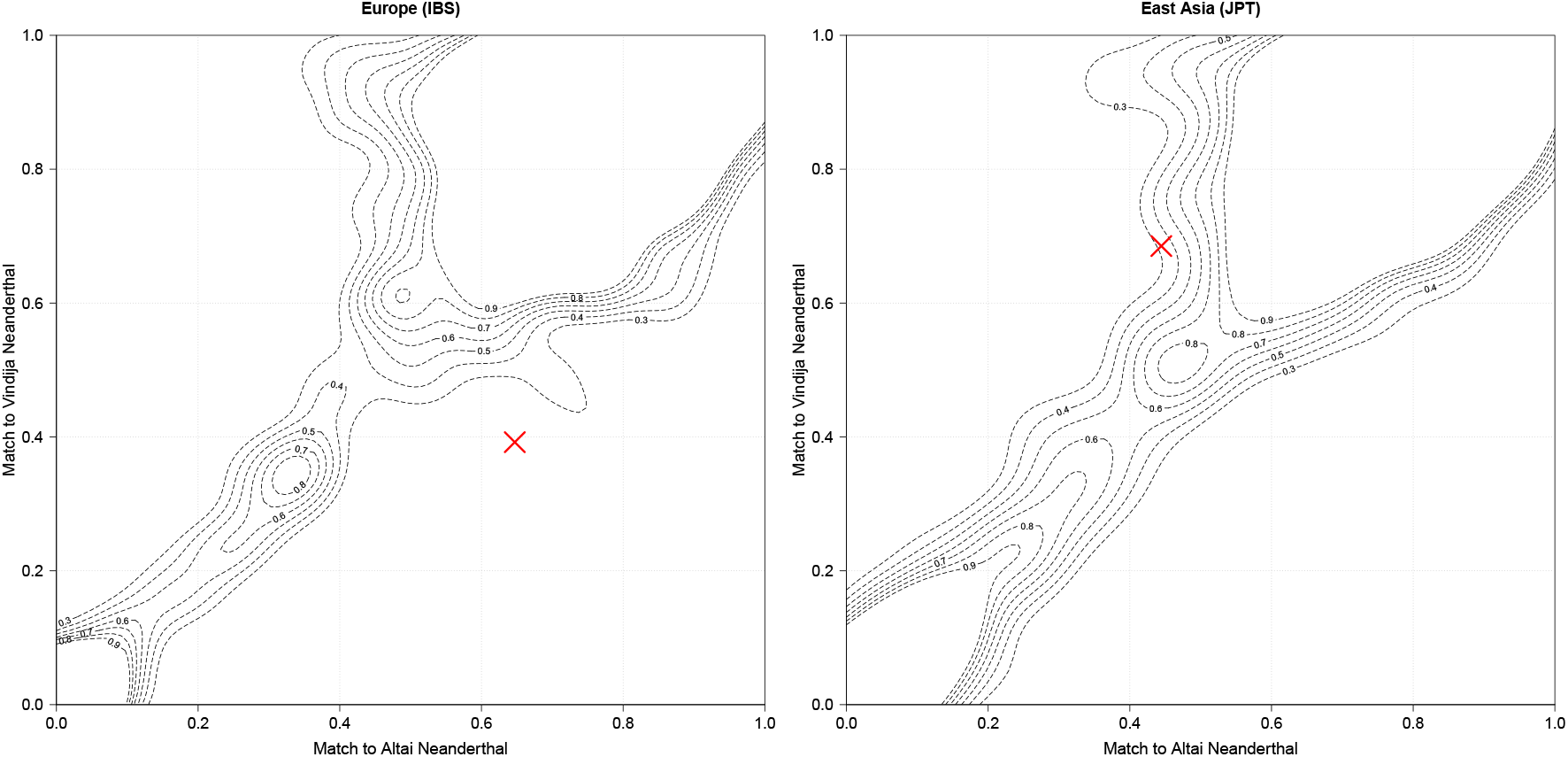
Contour density plots of match proportion of human-Neanderthal introgressed segments to the actual Vindija and Altai Neanderthal genomes. The plot visualizes the affinity of all Sprime inferred fragments to the Altai and Vindija Neanderthals (0=completely different, 1=identical to Neanderthal). The red x marks the human-Neanderthal introgressed ABO alleles found in a) European and b) Asian populations.

We plotted the introgressed fragments detected by Sprime to visualize them on an x-y axis. If all modern human alleles in an introgressed fragment are identical to the version in either Neanderthal chromosome, the match score is 1. If none of the alleles match, the score is 0. Scores in between reflect the proportion of alleles that match over all alleles. As observed in the figure, a large number of fragments match both Neanderthals highly, noted by the high density of fragments at the top right corner. Conversely, fragments that have very low affinities to the Neanderthals, plotted in the bottom left corner, represent either false positive fragment calls, or introgressed fragments that match to a different archaic species.

Our expectation is for the introgressed segments containing the ABO gene to be similar but not identical to the versions in the actual Neanderthal genomes. We expect this because the Neanderthal population which introgressed with modern humans did not include the Vindija or Altai individuals, and thus there is expected sequence divergence between these three Neanderthal populations, reflected in an intermediate affinity to both Altai and Vindija individuals.

Our plot confirms that the European/Southeast Asian introgressed O2 haplotype has a high affinity to the Altai Neanderthal (¿0.6), but conversely, its affinity to the Vindija Neanderthal is lower than other genome fragments (¡0.4), falling outside of the majority of genome fragments. The East Asian introgressed O1 haplotype has a high affinity to the Vindija Neanderthal (¿0.6), and a somewhat low affinity to the Altai Neanderthal (¿0.4). Neither introgressed haplotype was identical to the actual Neanderthal version.

The asymmetry in higher affinities to one but not both Neanderthals makes the introgressed ABO haplotypes fall outside the variation of other introgressed genome fragments in Europe, and on the margins of other introgressed genome fragments in East Asia. This pattern suggests that the Neanderthal population that introgressed with modern humans carried ABO haplotypes that were diverse relative to all other introgressed genome fragments.

## 4 Discussion

We found that archaic humans possessed unique ABO haplotypes, which are structurally similar to modern human ABO alleles, and in some cases are still found in modern humans through introgression. The similarities found in the coding region suggest those alleles are functionally identical to modern ABO alleles. While it is difficult to speculate on the selective background for archaic specific ABO haplotypes, it is interesting to find Neanderthal ABO alleles at a moderate frequencies in modern humans, as there is compelling evidence for strong selection against Neanderthal versions of functional genes [Sankararaman et al., 2016, Harris and Nielsen, 2016, Petr et al., 2019, Juric et al., 2016]. The most likely explanation is that both Neanderthal O alleles identified here are selectively neutral relative to modern human O alleles, and thus its frequency in modern humans is just a consequence of neutral demographic effects.

Finding a O1 allele in the Vindija individual from Croatia is consistent with a previous study by Lalueza-Fox et al. [2008], which found the deletion responsible for the common O1 and O1v alleles in two other European Neanderthals from El Sidron. Furthermore, we find the same O1 allele in the Vindija and the Chagyrskaya Neanderthal (from Siberia), and we find that both present similar genetic distances to introgressed alleles in modern populations (Figure 6). This is again consistent with the analysis of the Chagyrskaya Neanderthal genome in Mafessoni et al. [2020], in which Chagyrskaya is more closely related to European Neanderthals than to other Siberian Neanderthals such as Altai.

The geographic distribution of Neanderthal O alleles in modern human populations; where the Altai/Chagyrskaya O2 type is found in Europe and Southeast Asia, and the Vindija/Chagyrskaya O1 type is found exclusively in East Asia, is consistent with independent Neanderthal-modern human introgression events. This result is consistent with findings in other genomic regions, where at least two Neanderthal variants are found in modern humans with different geographic distributions [Taskent et al., 2020, Zeberg et al., 2020]. However, it is possible that both Neanderthal O alleles were maintained in the same Neanderthal population or populations that interbred with modern humans, and these haplotypes segregated in modern human populations due to diverse demographic forces between European, Southeast Asian, and East Asian populations. It is important to remember here, that neither Vindija or Altai were part of the Neanderthal population that directly interbred with modern humans, but were instead represent three different Neanderthal lineages [Prüfer et al., 2017].

Finally, based on our affinity plot, we see that all introgressed genome fragments found in the European and East Asian individuals present similar affinity to both Neanderthal individuals (following a diagonal line in the affinity map), with a skew towards the Vindija Neanderthal, as this individual is more closely related to the population that interbred with modern humans than the Altai Neanderthal [Prüfer et al., 2017].

Unexpectedly, this is not the pattern we observe for the genome fragment that contains the ABO gene, instead for each population, these fragments have an asymmetric affinity for only one Neanderthal individual. We find that the modern Altai/Chagyrskaya-like O2 allele found in Europeans has a higher affinity for the Altai individual than the Vindija individual, and appears as an outlier in the relative affinity of all other genome fragments (Figure 5a). Similarly, the Vindija/Chagyrskaya-like introgressed O1 allele found in East Asians has a higher affinity for the Vindija individual than the Altai individual, and its affinity falls on the edge of the relative affinity of all other genome fragments (Figure 5b).

The similar affinities to both individuals we observe in all other introgressed fragments suggests shared ancestry between the introgressing Neanderthal population, the Altai population, and the Vindija population. ABO is an exception, suggesting Neanderthals are more diverse at the ABO locus than other regions of the genome. Given that neither the Vindija or Altai individuals were part of the population that directly interbred with modern humans, the most consistent explanation for this pattern is that these two alleles were maintained in the Neanderthal population or populations that interbred with modern humans, retaining a larger haplotype diversity relative to other genome elements. If this was the case, these various haplotypes were likely retained in different Neanderthal populations by balancing selection, which is expected to maintain ancestral diversity much longer than neutral regions of the genome, just as balancing selection maintains ABO variation in modern humans.

### 4.1 Incomplete Lineage Sorting

A potential limitation of this study is the effect of incomplete lineage sorting (ILS), in which ancestral ABO haplotypes could be retained in both Neanderthals and modern human lineages since the divergence of both species. These ancestrally retained haplotypes would then mimic true human-Neanderthal introgression.

While the long divergence time between modern humans and Neanderthals makes this possibility unlikely, as recombination over time would break up these haplotypes. We have calculated the probability of the genome fragment carrying the ABO gene to be maintained in both human and Neanderthal lineages through ILS to very small (p = 5.5e-15), further supporting that the Neanderthal-like O alleles found in modern humans to be the result of admixture between the two species (for details, see the Supplement).

### 4.2 Conclusion

Human genetic variation in the ABO gene is a classic marker for genetic diversity in humans [Cavalli-Sforza et al., 1964]. Here, we provide an in-depth description of the genetic diversity of the ABO gene in four archaic humans, based on published ancient genomes. We found that archaic ABO haplotypes are polymorphic at the same positions which define modern human ABO function, and have posited that these archaic alleles must function similarly to modern human alleles. Furthermore, we found two Denisovan-specific O haplotypes, which are genetically similar to modern O alleles, while Neanderthal-specific O haplotypes are derived relative to human alleles, but found today at low frequencies due to past human-Neanderthal admixture.

Finding four different Neanderthal variants in late-era Neanderthals (*<*120,000 YBP) is unexpected. A common perception, based on long runs of homozygosity seen across Neanderthal genomes, is that late-era Neanderthals were extremely inbred and thus had reduced genetic diversity. The high allele diversity found in these Neanderthals was possibly maintained through balancing selection at the ABO locus. This notion dovetails with our contour map results, showing introgressed Neanderthal O haplotypes falling outside of the genome divergence of all other fragments, and suggests that balancing selection operated in Neanderthals similarly to modern humans.

## 5 Acknowledgements

The authors would like to thank Alyssa Funk, for checking the ABO gene genome coordinates against ancestry genome tracks for admixed individuals in the 1000 Genomes Project panel, Dr. Sharon Browning, for providing previously unpublished data on Vindija genome affinity to the populations in the 1000 Genomes Project, and Dr. Kelsey Witt and Dr. Iain Mathieson for comments on early versions of this manuscript. FAV was supported by NIH grant R35GM128946 (to EHS).

## 6 Methods

### 6.1 Haplotype construction and determination of ABO subtype from NGS datasets

Coding sequence data for the ABO locus (5,008 ABO chromosomes) were obtained from a publicly available global reference panel, the 1,000 Genomes Project (Phase III), which contains a diverse set of individuals from multiple populations [Sudmant et al., 2010]. We obtained genotype calls for the ABO locus from each NGS dataset included in our analysis in variant call format (VCF) file using the Genome Analysis Tool Kit (GATK) to call single nucleotide variants (SNVs) as well as insertions and deletions (indels) [McKenna et al., 2010]. Both indels and SNVs are important for blood group calling because the primary differences between the A and B haplotypes are SNVs while the common cause of the O blood type is a single base deletion that causes nonsense mediated decay of the RNA transcript resulting in absence of protein [Yamamoto et al., 2012]. In order to resolve ABO haplotypes from these NGS datasets, we employed PHASE 2.1.2 for haplotype construction of the different chromosomal alleles for each individual [Scheet and Stephens, 2006]. The 68 sites used to determine ABO alleles are described in Yip [2002], Patnaik et al. [2012]. A comprehensive summary of methods used to call blood group variants from NGS data can be found in Wheeler and Johnsen [2018].

### 6.2 Inclusion of high coverage paleogenomic datasets

In addition to analyzing ABO haplotype structure in the 1,000 Genomes Project panel, we extracted coding variation from four archaic human genomes; pertaining to a Neanderthal individual from Croatia (∼42X coverage for the coding region of ABO) [Prüfer et al., 2017], two Neanderthal individuals from the Altai Mountains: one from the Denisova cave (∼30X coverage for the coding region of ABO) [Prüfer et al., 2014], and one from the Chagyrskaya cave (∼28x coverage for the coding region of ABO) [Mafessoni et al., 2020]. Finally, a single Denisovan individual from the Denisova cave in the Altai Mountains (∼21X coverage for the coding region of ABO) [Meyer et al., 2012]. These individuals are estimated to be at least 50,000 years old.

To analyse the genetic distance between archaic and modern ABO haplotypes, we used sequence data from VCF files for all archaic individuals. We use the complete haplotype for the ABO gene (Hg19 position 9:136130563-136150630), which contains unphased exonic and intronic data. To then impute function from these sequences, we extracted all 68 variable loci identified in our initial haplotype construction of the 1,000 Genomes Project, and characterized them in all archaic humans by calling variation from the alignment of archaic raw reads in BAM format.

### 6.3 Haplotype distances

We used Haplostrips [Marnetto and Huerta-Sánchez, 2017] to quantify the relatedness of modern ABO haplotypes with the archaic haplotypes from the Neanderthal and Denisovan individuals. For this analysis, we use the complete haplotype for the ABO gene (Hg19 position 9:136130563-136150630), which contains exonic and intronic data, found in the variant call format (VCF) files from the 1000 Genomes Project and the various archaic genome projects. Each haplostrip is polarized to one the archaic genome (reference haplotype), and each subsequent haplotype is ordered by genetic similarity, from most related to least related. The archaic haplotypes are unphased (only coding regions were phased for calling functional ABO alleles). For this analysis, heterozygous positions are assigned between the two chromosomes randomly. Although not phased, both archaic chromosomes share a large amount of unique polymorphism reflecting the separate evolutionary history of archaic species, and thus will cluster together, pulling other introgressed archaic haplotypes closer than other non-introgressed haplotypes, as those haplotypes were also shaped by the evolutionary history of the archaic species.

We used the genetic distances calculated by Haplostrips to rank the proximity of modern human haplotypes to archaic haplotypes. We used R [R Development Core Team, 2008] to visualize the genetic distances between individuals in the 1000 genomes populations to the archaic ABO haplotypes. The distance information we used to generate these figures can be found in the supplement.

Identifying the origin of archaic introgression in admixed populations is complex. For example, American populations can trace portions of their genomes to European and African ancestry, as a consequence of European colonization of Native Americans, and the African slave trade. Because of these historical events, archaic introgression present in modern American individuals can be inherited from any of these sources. For the American populations in the 1000 Genomes Project panel, including two African populations sampled in the Southwest United States and Barbados, we used ancestry calls from Martin et al. [2017] to distinguish if purported archaic ABO alleles sit in European-ancestry genome tracks. This allowed us to properly track archaic introgression to Europe, rather than being retained ancestrally in African populations, or introgression with the Asian ancestral populations from which Native Americans descend. For our genetic distance results, we thus excluded six populations sampled in America (MXL, PUR, PEL, CLM), including two African populations (ASW,ACB), after determining that all introgresssed ABO haplotypes are exclusively located in European ancestry tracks, thus providing a confusing look at archaic introgression (see Supp. Table 4).

### 6.4 Imputing functional ABO alleles from the archaic data

To impute function from the archaic data, we extracted the 68 polymorphic positions that define ABO allele variation, identified from modern humans in the previous section. We then used ANGSD [Korneliussen et al., 2014] to count the number of each base pair (A,C,G,T) at each position. In order to distinguish true polymorphism at heterozygous sites from sequencing errors and aDNA damage, we only considered base pairs that are represented more than two times at each position in the BAM read alignment. Because DNA damage and sequencing error is distributed randomly, we expect false polymorphism to only appear once at each position, but we removed bases appearing up to twice for redundancy. Base pair counts, genotypes, and coverage depth can be found at: https://drive.google.com/drive/folders/1a6ct0nYc6aCupWUWmhTh5FAfQcG1x9qF?usp=sharing. Any remaining polymorphism was evaluated by observing the alignment directly using Geneious Prime 2020.2.4 (https://www.geneious.com). Screenshots of polymorphic sites can be found at: https://drive.google.com/drive/folders/1F1s4xXO5TetYSJJWeFEnakOhNCjK3kW1?usp=sharing.

Understanding the function of any archaic gene is limited by our ability to phase archaic genome sequences, as population-based phasing methods cannot be applied directly to a single individual from an extinct population [Browning and Browning, 2011]. In our study, we mitigate the impact of incorrect phasing in the following ways. First, we used the raw reads to link any two polymorphic sites that are shared in a single read (Supp. Figure 3). Then, for all unlinked blocks, we explored all possible combinations by looking for a match for the 68 variable sites in modern humans ABO haplotypes. We find that several combinations are not represented in modern humans, and use this as the first line of evidence to assign phasing (Supp. Figure 4).

Second, we attempt to use archaic genome information (without using modern information) to assign phasing. This is possible because the Altai Neanderthal is phased (homozygous at all positions) and presents two identical O2 alleles. We then use the Altai O2 allele to phase the three heterozygous blocks in the Chagyrskaya Neanderthal, by assuming both share a O2 allele. This results in the Chagyrskaya Neanderthal presenting an O2 and an O1 alleles. We then use the Chagyrskaya O1 allele to attempt to phase the four heterozygous blocks in the Vindija Neanderthal, assuming that both share an O1 allele. This results in the Vindija Neanderthal presenting an O1 allele, and a cis-AB (B(A)) allele, which is consistent with the first heuristic we used to solve phasing (Supp. Figure 5).

Third, we take advantage of identifying introgressed Vindija-like O1 and Altai-like O2 alleles in modern humans. These are Neanderthal ABO sequences that survive in modern humans, which we use as a template to phase heterozygous positions to either chromosome (Supp. Figure 6). This phasing results in the Chagyrskaya Neanderthal presenting an O2 and an O1 alleles, and the Vindija Neanderthal presenting an O1 allele, and a cis-AB (B(A)) allele, which is consistent with the first and second heuristic we used to solve phasing. A table with 68 variable positions including variation for the phased archaic haplotypes can be found at: https://drive.google.com/file/d/1diIy7mZUH32-X299VRhJzQJ7GgvLR77n/view?usp=sharing.

### 6.5 Contour density plots of match proportion of introgressed segments to the Neanderthal genomes

In order to look more closely at the Neanderthal ABO alleles in present-day humans, we plotted two-way densities of match rate to the Altai Neanderthal and Vindija Neanderthal genomes, using the method described in Browning et al. [2018]. We used the archaic SNP calls generated in Browning et al. [2018] using Sprime statistics. We also used Browning et al. [2018]’s match-mismatch calls to check if the purported introgressed SNPs also match the Vindija, and Altai archaic genomes. We used R [R Development Core Team, 2008] to visualize the affinity data. We plotted a contour density figure using the Neanderthal match/non-match scores in the Iberian (IBS) and Japanese (JPT) 1000 Genomes Panel populations, reported in Browning et al. [2018]. We mapped the affinity of all introgressed genome fragments to either the Altai and Vindija genomes in these two representative European and East Asian populations. We then matched the variable sites in the sequence of the Altai-like and Vindija-like introgressed ABO haplotypes to the Altai and Vindija genomes to visualize their affinity relative to other introgressed genome fragments. The resulting contour density plots is interpreted as a topological map, showing the affinity of all introgressed fragments in a modern population relative to the two Neanderthal genomes. Then, we highlight the position of the introgressed ABO haplotypes relative to all other introgressed genome element, in order to quantify their relative sequence divergence.

## References

Sharon R Browning and Brian L Browning. Haplotype phasing: existing methods and new developments. Nature Reviews Genetics, 12(10):703, 2011.

Sharon R Browning, Brian L Browning, Ying Zhou, Serena Tucci, and Joshua M Akey. Analysis of human sequence data reveals two pulses of archaic denisovan admixture. Cell, 173(1):53–61, 2018.

Luigi Luca Cavalli-Sforza, I Barrai, and AWF Edwards. Analysis of human evolution under random genetic drift. In Cold Spring Harbor symposia on quantitative biology, volume 29, pages 9–20. Cold Spring Harbor Laboratory Press, 1964.

Sejong Chun, Sooin Choi, HongBi Yu, and Duck Cho. Cis-ab the blood group of many faces, is a conundrum to the novice eye. Annals of laboratory medicine, 39(2):115–120, 2019.

Benito Estrada-Mena, F Javier Estrada, Raúl Ulloa-Arvizu, Miriam Guido, Rocío Méndez, Ramón Coral, Thelma Canto, Julio Granados, Rodrigo Rubí-Castellanos, Héctor Rangel-Villalobos, et al. Blood group o alleles in native americans: implications in the peopling of the americas. American Journal of Physical Anthropology: The Official Publication of the American Association of Physical Anthropologists, 142(1):85–94, 2010.

Andrew E Fry, Michael J Griffiths, Sarah Auburn, Mahamadou Diakite, Julian T Forton, Angela Green, Anna Richardson, Jonathan Wilson, Muminatou Jallow, Fatou Sisay-Joof, et al. Common variation in the abo glycosyltransferase is associated with susceptibility to severe plasmodium falciparum malaria. Human molecular genetics, 17(4):567–576, 2007.

Richard E Green, Johannes Krause, Adrian W Briggs, Tomislav Maricic, Udo Stenzel, Martin Kircher, Nick Patterson, Heng Li, Weiwei Zhai, Markus Hsi-Yang Fritz, et al. A draft sequence of the neandertal genome. science, 328(5979):710–722, 2010.

Kelley Harris and Rasmus Nielsen. The genetic cost of neanderthal introgression. Genetics, 203(2):881–891, 2016.

Emilia Huerta-Sánchez and Fergal P Casey. Archaic inheritance: supporting high-altitude life in tibet. Journal of Applied Physiology, 119(10):1129–1134, 2015.

Emilia Huerta-Sánchez, Xin Jin, Zhuoma Bianba, Benjamin M Peter, Nicolas Vinckenbosch, Yu Liang, Xin Yi, Mingze He, Mehmet Somel, Peixiang Ni, et al. Altitude adaptation in tibetans caused by introgression of denisovan-like dna. Nature, 512(7513): 194, 2014.

Ivan Juric, Simon Aeschbacher, and Graham Coop. The strength of selection against neanderthal introgression. PLoS genetics, 12(11):e1006340, 2016.

Thorfinn S. Korneliussen, Anders Albrechtsen, and Rasmus Nielsen. ANGSD: Analysis of next generation sequencing data. BMC Bioinformatics, 15(1):356, November 2014. ISSN 1471-2105. doi: 10.1186/s12859-014-0356-4. URL http://www.biomedcentral.com/1471-2105/15/356/abstract.

Carles Lalueza-Fox, Elena Gigli, Marco de la Rasilla, Javier Fortea, Antonio Rosas, Jaume Bertranpetit, and Johannes Krause. Genetic characterization of the abo blood group in neandertals. BMC evolutionary biology, 8(1):342, 2008.

Fabrizio Mafessoni, Steffi Grote, Cesare de Filippo, Viviane Slon, Kseniya A. Kolobova, Bence Viola, Sergey V. Markin, Manjusha Chintalapati, Stephane Peyrégne, Laurits Skov, Pontus Skoglund, Andrey I. Krivoshapkin, Anatoly P. Derevianko, Matthias Meyer, Janet Kelso, Benjamin Peter, Kay Prüfer, and Svante Pääbo. A high-coverage neandertal genome from chagyrskaya cave. bioRxiv, 2020. doi: 10.1101/2020.03.12.988956. URL https://www.biorxiv.org/content/early/2020/03/13/2020.03.12.988956.

Davide Marnetto and Emilia Huerta-Sánchez. Haplostrips: revealing population structure through haplotype visualization. Methods in Ecology and Evolution, 8(10):1389–1392, 2017.

Alicia R Martin, Christopher R Gignoux, Raymond K Walters, Genevieve L Wojcik, Benjamin M Neale, Simon Gravel, Mark J Daly, Carlos D Bustamante, and Eimear E Kenny. Human demographic history impacts genetic risk prediction across diverse populations. The American Journal of Human Genetics, 100(4):635–649, 2017.

Aaron McKenna, Matthew Hanna, Eric Banks, Andrey Sivachenko, Kristian Cibulskis, Andrew Kernytsky, Kiran Garimella, David Altshuler, Stacey Gabriel, Mark Daly, et al. The genome analysis toolkit: a mapreduce framework for analyzing next-generation dna sequencing data. Genome research, 20(9):1297–1303, 2010.

Matthias Meyer, Martin Kircher, Marie-Theres Gansauge, Heng Li, Fernando Racimo, Swapan Mallick, Joshua G Schraiber, Flora Jay, Kay Prüfer, Cesare De Filippo, et al. A high-coverage genome sequence from an archaic denisovan individual. Science, 338 (6104):222–226, 2012.

Santosh Kumar Patnaik, Wolfgang Helmberg, and Olga O Blumenfeld. Bgmut: Ncbi dbrbc database of allelic variations of genes encoding antigens of blood group systems. Nucleic acids research, 40(D1):D1023–D1029, 2012.

Martin Petr, Svante Pääbo, Janet Kelso, and Benjamin Vernot. Limits of long-term selection against neandertal introgression. Proceedings of the National Academy of Sciences, 116(5):1639–1644, 2019.

Kay Prüfer, Fernando Racimo, Nick Patterson, Flora Jay, Sriram Sankararaman, Susanna Sawyer, Anja Heinze, Gabriel Renaud, Peter H Sudmant, Cesare De Filippo, et al. The complete genome sequence of a neanderthal from the altai mountains. Nature, 505(7481): 43, 2014.

Kay Prüfer, Cesare de Filippo, Steffi Grote, Fabrizio Mafessoni, Petra Korlević, Mateja Hajdinjak, Benjamin Vernot, Laurits Skov, Pinghsun Hsieh, Stéphane Peyrégne, et al. A high-coverage neandertal genome from vindija cave in croatia. Science, 358(6363): 655–658, 2017.

R Development Core Team. R: A Language and Environment for Statistical Computing. R Foundation for Statistical Computing, Vienna, Austria, 2008. URL http://www.R-project.org. ISBN 3-900051-07-0.

Fernando Racimo, Sriram Sankararaman, Rasmus Nielsen, and Emilia Huerta-Sánchez. Evidence for archaic adaptive introgression in humans. Nature Reviews Genetics, 16(6): 359, 2015.

Fernando Racimo, Davide Marnetto, and Emilia Huerta-Sanchez. Signatures of archaic adaptive introgression in present-day human populations. Molecular biology and evolution, 34(2):296–317, 2016.

Francis Roubinet, Stéphanie Despiau, Francesc Calafell, Fen Jin, Jaume Bertanpetit, Naruya Saitou, and Antoine Blancher. Evolution of the o alleles of the human abo blood group gene. Transfusion, 44(5):707–715, 2004.

Sriram Sankararaman, Swapan Mallick, Nick Patterson, and David Reich. The combined landscape of denisovan and neanderthal ancestry in present-day humans. Current Biology, 26(9):1241–1247, 2016.

Paul Scheet and Matthew Stephens. A fast and flexible statistical model for large-scale population genotype data: applications to inferring missing genotypes and haplotypic phase. The American Journal of Human Genetics, 78(4):629–644, 2006.

Laure Ségurel, Emma E Thompson, Timothée Flutre, Jessica Lovstad, Aarti Venkat, Susan W Margulis, Jill Moyse, Steve Ross, Kathryn Gamble, Guy Sella, et al. The abo blood group is a trans-species polymorphism in primates. Proceedings of the National Academy of Sciences, 109(45):18493–18498, 2012.

Robert M Seymour, Martin J Allan, Andrew Pomiankowski, and Kenth Gustafsson. Evolution of the human abo polymorphism by two complementary selective pressures. Proceedings of the Royal Society of London. Series B: Biological Sciences, 271(1543):1065–1072, 2004.

Peter H Sudmant, Jacob O Kitzman, Francesca Antonacci, Can Alkan, Maika Malig, Anya Tsalenko, Nick Sampas, Laurakay Bruhn, Jay Shendure, Evan E Eichler, et al. Diversity of human copy number variation and multicopy genes. Science, 330(6004):641–646, 2010.

Ozgur Taskent, Yen Lung Lin, Ioannis Patramanis, Pavlos Pavlidis, and Omer Gokcumen. Analysis of haplotypic variation and deletion polymorphisms point to multiple archaic introgression events, including from altai neanderthal lineage. Genetics, 2020.

Fernando A Villanea and Joshua G Schraiber. Multiple episodes of interbreeding between neanderthal and modern humans. Nature ecology & evolution, 3(1):39, 2019.

Fernando A Villanea, Deborah A Bolnick, Cara Monroe, Rosita Worl, Rosemary Cambra, Alan Leventhal, and Brian M Kemp. Brief communication: Evolution of a specific o allele (o1vg542a) supports unique ancestry of native americans. American journal of physical anthropology, 151(4):649–657, 2013.

Fernando A Villanea, Kristin N Safi, and Jeremiah W Busch. A general model of negative frequency dependent selection explains global patterns of human abo polymorphism. PloS one, 10(5):e0125003, 2015.

Marsha M Wheeler and Jill M Johnsen. The role of genomics in transfusion medicine. Current opinion in hematology, 25(6):509–515, 2018.

Fumi-ichiro Yamamoto, Patricia D McNeill, and Sen-itiroh Hakomori. Human histo-blood group a2 transferase coded by a2 allele, one of the a subtypes, is characterized by a single base deletion in the coding sequence, which results in an additional domain at the carboxyl terminal. Biochemical and biophysical research communications, 187(1): 366–374, 1992.

Fumiichiro Yamamoto, Emili Cid, Miyako Yamamoto, and Antoine Blancher. Abo research in the modern era of genomics. Transfusion medicine reviews, 26(2):103–118, 2012.

SP Yip. Sequence variation at the human abo locus. Annals of human genetics, 66(1): 1–27, 2002.

Hugo Zeberg, Janet Kelso, and Svante Pääbo. The neandertal progesterone receptor. Molecular Biology and Evolution, 2020.

Xinjun Zhang, Bernard Kim, Kirk E Lohmueller, and Emilia Huerta-Sánchez. The impact of recessive deleterious variation on signals of adaptive introgression in human populations. Genetics, 215(3):799–812, 2020.

