## Supplementary Materials for "ABO genetic variation in Neanderthals and Denisovans"

Compiled on December 24, 2020.

#### 1 Sharing of Neanderthal ABO haplotypes through Incomplete Lineage Sorting

We calculate the probability of a genome fragment carrying the ABO gene, of length of 31kb, shared by modern humans and Neanderthals due to incomplete ancestral lineage sorting, as described in Huerta-Sánchez et al. [2014]. For this calculation, r is the recombination rate per generation per bp, of 3.39e-8, for a genome fragment at coordinates HG19 9:136125329-136157138, as described in the HapMap recombination map [Consortium et al., 2007] and found at: `ftp://ftp-trace.ncbi.nih.gov/1000genomes/ftp/technical/working/20110106_recombination_hotspots/`.

The divergence time of the human and Altai Neanderthal branches t, is 980,000 years, which is calculated as twice the split between Neanderthals and modern humans (550,000 years) minus the time at sampling of the Altai Neanderthal (120,000 years) [Prüfer et al., 2014, 2017, Douka et al., 2019]. We used an estimated time for interbreeding between the two groups of 50,000 years ago, and a generation time of 29 years. Under these assumptions, we exclude that it derives from the common ancestor (p = 5.5e-15) and conclude that this region entered the human gene pool through admixture with Neanderthals. Furthermore, we recalculated this probability using an extremely conservative divergence time of 300,000 years, as used for Denisovans in Huerta-Sánchez et al. [2014]. We should mention, using a highly conservative Neanderthalhuman split time considerably increases the probability of incomplete lineage sorting. Using this extreme estimate, the probability of incomplete lineage sorting is still only p = 0.00017, further supporting that this region entered the human gene pool through admixture with Neanderthals.

| Superpopulation | Population | Fragment ID | Chromosome 9 Position |
|---|---|---|---|
| Europe | GBR | 15 | 136071592-136374869 |
| Europe | IBS | 37 | 136109740-136412483 |
| Europe | TSI | 24 | 136071592-136383617 |
| Southeast Asia | BEB | 18 | 135862746-136374869 |
| Southeast Asia | GIH | 23 | 136131316-136374869 |
| Southeast Asia | ITU | 15 | 136077559-136389868 |
| East Asia | JPT | 29 | 136131539-136368328 |

Table 1: Superpopulation, Population, ID, and hg19 coordinates for archaic genome fragments in the 1000 Genomes Project detected in Browning et al. [2018] which contain the ABO gene.

#### 2 Supplementary Tables

Supplementary Table 1. Superpopulation, Population, ID, and hg19 coordinates for archaic genome fragments in the 1000 Genomes Project detected in Browning et al. [2018] which contain the ABO gene.

#### 3 Admixed populations in the 1000 Genomes Panel

Supplementary Table 2. Chromosome, start and stop position, ancestry block (NAT=Native American, EUR=European, AFR=African),for all neanderthal introgressed ABO haplotypes in American individuals, identified by a sample ID, and 1000 Genomes Panel population. In all individuals the neanderthal introgressed haplotype was found in an EUR ancestry block, confirming that these haplotypes were inherited through European admixture post-American colonization. Find .txt file at: `https://drive.google.com/file/d/1ckuFOMZJ-0vmlofh_CqB1_Z1yNgsuU5P/view?usp=sharing`

#### 4 Haplostrips genetic distances (unsorted)

##### 4.1 Denisovan

Find .txt file at: `https://drive.google.com/file/d/1gKW9jQo7Sw0ycfnHG1BDgEql5r_X1AVr/view?usp=sharing`

##### 4.2 Altai neanderthal

Find .txt file at: `https://drive.google.com/file/d/1OgpVr99xNBC0LbqJ4ZQYxzVLyl4e5oX-/view?usp=sharing`

##### 4.3 Vindija neanderthal

Find .txt file at: `https://drive.google.com/file/d/1-gmqPSS35wiBYpBkGS6GCvEhIvCAc9z5/view?usp=sharing`

##### 4.4 Chagyrskaya neanderthal

Find .txt file at: `https://drive.google.com/file/d/1XFej1HItNtrN13m1H3dJeWPbxrTSn1D-/view?usp=sharing`

### 5 Supplementary Figures

Supplementary Figure 1

Figure allele variant table.pdf

| DBSNP | position | reference | sample | functionGVS | rsID | amino acid | protein | polyPhen | GERP | ExAC |
|---|---|---|---|---|---|---|---|---|---|---|
| dbSNP_129 | 136131056 | CG | C | frameshift | 56392308 | none | NA | unknown | -5.32 | delG=5059/ref=40137 |
| none | 136131059 | G | G/C | missense | 0 | ASN,LYS | 353/355 | 0.923 | -0.105 | C=1/G=49241 |
| none | 136131060 | T | T/G | missense | 0 | ASN,THR | 353/355 | 0.597 | -1.81 | G=8/T=49500 |
| dbSNP_129 | 136131064 | G | G/A | missense | 56390333 | ARG,TRP | 352/355 | 1 | 1.03 | C=2/A=41/G=54483 |
| dbSNP_135 | 136131065 | G | G/A | synonymous | 183748371 | none | 351/355 | unknown | -0.912 | A=3/G=56545 |
| dbSNP_116 | 136131069 | G | G/A | missense | 7466899 | ALA,VAL | 350/355 | 0.095 | -2.65 | A=78/G=62032 |
| none | 136131086 | C | C/T | synonymous | 0 | none | 344/355 | unknown | 3.47 | T=17/C=85651 |
| dbSNP_129 | 136131109 | T | T/C | missense | 56231718 | ARG,GLY | 337/355 | 0.47 | -0.002 | C=29/T=109011 |
| dbSNP_138 | 136131118 | C | C/T | missense | 369695939 | ALA,THR | 334/355 | 0.013 | -8.76 | T=2/C=113272 |
| none | 136131154 | C | C/T | missense | 0 | GLU,LYS | 322/355 | 1 | 4.38 | T=5/C=119169 |
| dbSNP_117 | 136131188 | C | C/T | synonymous | 8176749 | none | 310/355 | unknown | 1.36 | T=14613/C=106423 |
| dbSNP_129 | 136131192 | T | T/C | missense | 56346931 | TYR,CYS | 309/355 | 1 | 3.19 | C=115/T=120953 |
| none | 136131240 | G | G/C | missense | 0 | ALA,GLY | 293/355 | 0.022 | 0.545 | C=1/G=121047 |
| dbSNP_117 | 136131289 | C | C/T | missense | 8176748 | VAL,MET | 277/355 | 1 | 4.38 | A=1/T=31193/C=87126 |
| dbSNP_117 | 136131315 | C | C/G | missense | 8176747 | GLY,ALA | 268/355 | 0.006 | -8.39 | G=14032/C=93378 |
| dbSNP_127 | 136131316 | C | C/T | missense | 41302905 | GLY,ARG | 268/355 | 0.99 | 3.28 | T=1726/C=105746 |
| none | 136131319 | C | C/T | missense | 0 | GLY,ARG | 267/355 | 1 | 4.38 | G=1/T=1/C=106254 |
| dbSNP_117 | 136131322 | G | G/T | missense | 8176746 | LEU,MET | 266/355 | 0.045 | -8.76 | A=1/T=13903/G=91164 |
| dbSNP_117 | 136131347 | G | A/G | synonymous | 8176745 | none | 257/355 | unknown | -1.59 | A=25722/G=61360 |
| dbSNP_117 | 136131350 | G | G/T | synonymous | 8176744 | none | 256/355 | unknown | -3.34 | T=2927/G=70809 |
| none | 136131375 | C | C/G | missense | 0 | ARG,PRO | 248/355 | 1 | 1.42 | T=2/G=7/C=57099 |
| dbSNP_126 | 136131389 | G | G/A | synonymous | 35494115 | none | 243/355 | unknown | 3.48 | A=26/G=53258 |
| dbSNP_137 | 136131407 | G | G/A | synonymous | 201439325 | none | 237/355 | unknown | -0.929 | A=48/G=50292 |
| dbSNP_117 | 136131415 | C | C/T | missense | 8176743 | GLY,SER | 235/355 | 0.858 | 2.51 | T=8378/C=44320 |
| dbSNP_129 | 136131429 | C | C/T | missense | 56116432 | GLY,ASP | 230/355 | 1 | 4.39 | T=223/C=50585 |
| dbSNP_117 | 136131437 | C | C/T | synonymous | 8176742 | none | 227/355 | unknown | -8.65 | T=13460/C=43502 |
| dbSNP_117 | 136131461 | G | A/G | synonymous | 8176741 | none | 219/355 | unknown | -4.2 | A=9491/G=50041 |
| dbSNP_129 | 136131469 | G | G/A | missense | 56408700 | ARG,CYS | 217/355 | 0.037 | -9.38 | A=86/G=58342 |
| dbSNP_117 | 136131472 | A | A/T | missense | 8176740 | PHE,ILE | 216/355 | 1 | 3.55 | T=16226/A=47766 |
| dbSNP_135 | 136131490 | C | C/T | missense | 181536132 | VAL,MET | 210/355 | 0.934 | -9.38 | T=1/C=62309 |
| dbSNP_117 | 136131523 | G | G/A | missense | 8176739 | ARG,CYS | 199/355 | 0.868 | 2.8 | A=1405/G=66839 |
| dbSNP_129 | 136131539 | A | A/G | synonymous | 55764262 | none | 193/355 | unknown | 3.79 | G=74/A=77166 |
| dbSNP_135 | 136131556 | G | G/T | missense | 184446112 | ARG,SER | 188/355 | 1 | 2.75 | unknown |
| dbSNP_129 | 136131576 | C | C/T | stop-gained | 55727303 | TRP,stop | 181/355 | unknown | 4.56 | T=3278/C=98528 |
| dbSNP_129 | 136131589 | C | C/T | missense | 55687199 | ALA,THR | 177/355 | 0.048 | -6.69 | T=96/C=107234 |
| dbSNP_138 | 136131590 | G | G/A | synonymous | 371569951 | none | 176/355 | unknown | -9.38 | A=9/G=108075 |
| dbSNP_129 | 136131591 | C | C/T | missense | 56039827 | ARG,HIS | 176/355 | 0.024 | -1.05 | T=143/C=108359 |
| dbSNP_116 | 136131592 | G | C/G | missense | 7853989 | ARG,GLY | 176/355 | 0.011 | -1.09 | A=1/C=15935/G=94822 |
| none | 136131595 | C | C/T | missense | 0 | VAL,MET | 175/355 | 1 | 3.77 | T=51/C=112005 |
| dbSNP_129 | 136131616 | G | G/C | missense | 56043861 | ARG,GLY | 168/355 | 1 | -4.42 | A=1/C=2/G=115095 |
| dbSNP_129 | 136131621 | GT | G | frameshift | 56284703 | none | NA | unknown | -8.32 | delT=410/ref=115308 |
| dbSNP_129 | 136131630 | G | G/A | missense | 55756402 | THR,MET | 163/355 | 0.896 | 0.193 | A=82/G=116240 |
| dbSNP_137 | 136131635 | G | G/A | synonymous | 200932155 | none | 161/355 | unknown | -3.15 | A=139/G=116133 |
| dbSNP_117 | 136131636 | C | C/T | missense | 8176738 | ARG,HIS | 161/355 | 0 | -9.38 | A=1/T=84/C=116231 |
| dbSNP_86 | 136131651 | G | G/A | missense | 1053878 | PRO,LEU | 156/355 | 0.95 | 3.77 | A=10390/G=106232 |
| dbSNP_129 | 136131664 | A | A/G | missense | 55687353 | PHE,LEU | 152/355 | 0.983 | 4.69 | unknown |
| none | 136131704 | C | C/T | synonymous | 0 | none | 138/355 | unknown | 4.95 | T=1/C=116333 |
| none | 136131718 | G | G/A | synonymous | 0 | none | 134/355 | unknown | 4.16 | A=2/G=116272 |
| none | 136132845 | A | A/G | missense | 0 | PHE,LEU | 109/355 | 1 | 4.33 | G=4/A=122326 |
| dbSNP_117 | 136132852 | G | G/A | synonymous | 8176721 | none | 106/355 | unknown | -3.12 | A=1601/G=120723 |
| dbSNP_135 | 136132853 | T | T/C | missense | 181412963 | ASN,SER | 106/355 | 1 | 4.33 | C=2/T=122322 |
| none | 136132864 | G | G/A | synonymous | 0 | none | 102/355 | unknown | 0.112 | A=1/G=122311 |
| dbSNP_117 | 136132873 | T | C/T | synonymous | 8176720 | none | 99/355 | unknown | -8.39 | C=48708/T=73496 |
| dbSNP_117 | 136132908 | T | TC | frameshift | 8176719 | none | NA | unknown | 4.2 | insC=45730/ref=75728 |
| none | 136133466 | AG | A | frameshift | 0 | none | NA | unknown | 2.04 | unknown |
| dbSNP_83 | 136133506 | A | G/A | missense | 512770 | SER,PRO | 74/355 | unknown | 2.04 | G=92871/A=29225 |
| none | 136135226 | G | G/C | missense | 0 | PRO,ARG | 67/355 | 0.02 | 1.3 | C=1/G=122329 |
| dbSNP_138 | 136135232 | G | G/A | missense | 375733196 | SER,LEU | 65/355 | 0 | -1.07 | A=10/G=122320 |
| dbSNP_129 | 136135236 | C | C/T | missense | 56335272 | VAL,ILE | 64/355 | 0.137 | 1.3 | unknown |
| dbSNP_83 | 136135237 | A | A/G | coding-unknown | 549443 | none | NA | unknown | -2.61 | G=90060/A=31798 |
| dbSNP_83 | 136135238 | T | C/T | missense | 549446 | HIS,ARG | 63/355 | unknown | -2.61 | unknown |
| none | 136136728 | C | C/G | missense | 0 | GLY,ARG | 50/355 | 0.004 | 0.822 | G=1/C=66129 |
| dbSNP_83 | 136136770 | A | A/C | missense | 688976 | PHE,VAL | 36/355 | unknown | -3.95 | C=66731/A=21321 |
| dbSNP_117 | 136136773 | C | C/T | missense | 8176696 | GLY,ARG | 35/355 | 0.806 | 2.04 | A=2/T=1262/C=58966 |
| dbSNP_129 | 136137547 | C | C/A | missense | 55876802 | ARG,LEU | 18/355 | 0.001 | -2.13 | T=1/A=1785/C=109688 |
| none | 136137551 | G | G/A | missense | 0 | LEU,PHE | 17/355 | 0.007 | 1.37 | A=3/G=109807 |
| dbSNP_129 | 136137554 | C | C/T | missense | 55917063 | ALA,THR | 16/355 | 0.062 | -4.6 | G=2/T=215/C=108557 |
| dbSNP_129 | 136137555 | G | G/A | synonymous | 61736301 | none | 15/355 | unknown | 1.36 | A=110/G=107902 |
| dbSNP_138 | 136150600 | G | G/A | synonymous | 367824410 | none | 2/355 | unknown | -1.44 | A=11/G=4317 |

Figure 1: SNVs and indels which define ABO allele variation in the coding portion of the ABO gene. These 68 variants were identified and annotated in Yip [2002], Patnaik et al. [2012].

### 6 Neanderthal haplotypes resolved

Supplementary Figure 2

figure phasing diagram.pdf

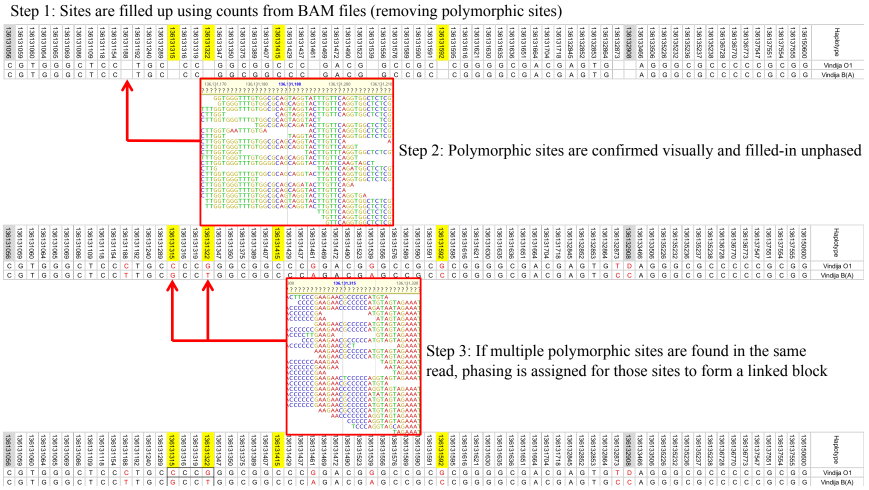


Figure 2: Steps taken to validate heterozyous sites, and link blocks of phased heterozygous sites

Supplementary Figure 3

Figure chagyrskaya phased.pdf

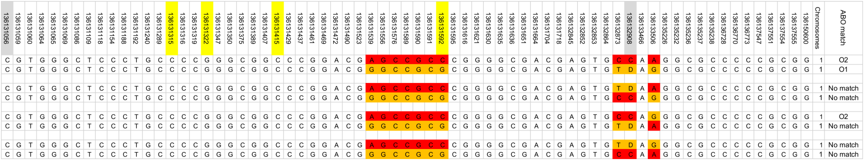

Figure 3: Possible configurations of unphased heterozygosity blocks for the two Chagyrskaya Neanderthal chromosomes, including closest match in modern ABO haplotypes

Supplementary Figure 4

Figure vindija phased.pdf

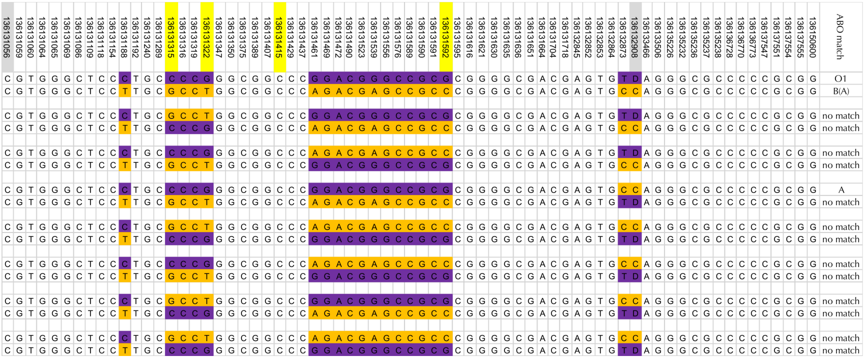

Figure 4: Possible configurations of unphased heterozygosity blocks for the two Vindija Neanderthal chromosomes, including closest match in modern ABO haplotypes

Supplementary Figure 5

Figure denisova phased.pdf

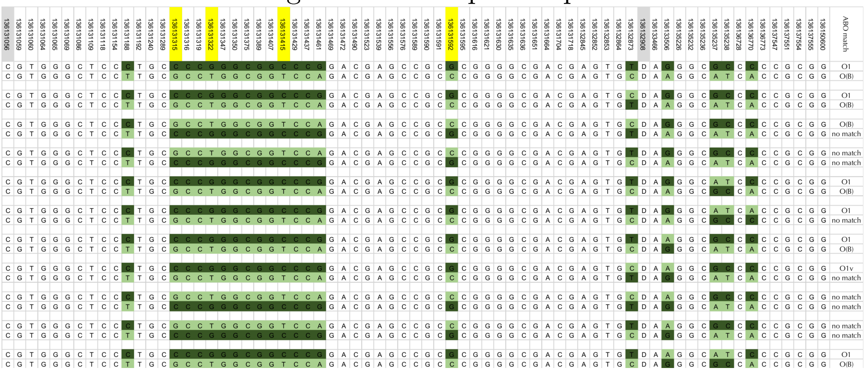

Figure 5: Possible configurations of unphased heterozygosity blocks for the two Denisovan chromosomes, including closest match in modern ABO haplotypes

#### 7 Introgressed genome fragments containing ABO

Supplementary Figure 6

Figure archaic aligned.pdf

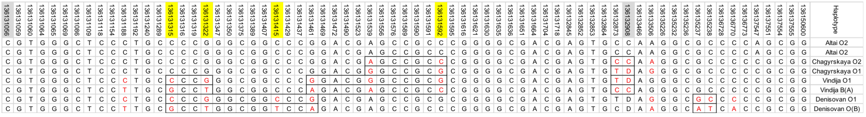

Figure 6: Phasing of ABO functional sites based on archaic variation only

Supplementary Figure 7

Figure surviving phasing.pdf

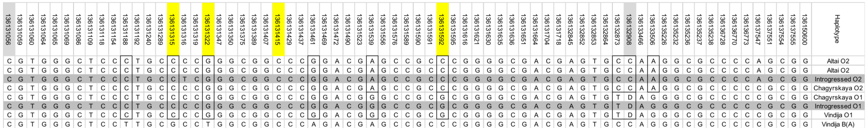

Figure 7: Phasing of ABO functional sites using surviving introgressed haplotypes as a reference

#### References


Sharon R Browning, Brian L Browning, Ying Zhou, Serena Tucci, and Joshua M Akey. Analysis of human sequence data reveals two pulses of archaic denisovan admixture. *Cell*, 173(1):53–61, 2018.

International HapMap Consortium et al. A second generation human haplotype map of over 3.1 million snps. *Nature*, 449(7164):851, 2007.

Katerina Douka, Viviane Slon, Zenobia Jacobs, Christopher Bronk Ramsey, Michael V Shunkov, Anatoly P Derevianko, Fabrizio Mafessoni, Maxim B Kozlikin, Bo Li, Rainer Grün, et al. Age estimates for hominin fossils and the onset of the upper palaeolithic at denisova cave. *Nature*, 565(7741): 640–644, 2019.

Emilia Huerta-Sánchez, Xin Jin, Zhuoma Bianba, Benjamin M Peter, Nicolas Vinckenbosch, Yu Liang, Xin Yi, Mingze He, Mehmet Somel, Peixiang Ni, et al. Altitude adaptation in tibetans caused by introgression of denisovan-like dna. *Nature*, 512(7513):194, 2014.

Santosh Kumar Patnaik, Wolfgang Helmberg, and Olga O Blumenfeld. Bgmut: Ncbi dbrbc database of allelic variations of genes encoding antigens of blood group systems. *Nucleic acids research*, 40(D1):D1023–D1029, 2012.

Kay Prüfer, Fernando Racimo, Nick Patterson, Flora Jay, Sriram Sankararaman, Susanna Sawyer, Anja Heinze, Gabriel Renaud, Peter H Sudmant, Cesare De Filippo, et al. The complete genome sequence of a neanderthal from the altai mountains. *Nature*, 505(7481):43, 2014.

Kay Prüfer, Cesare de Filippo, Steffi Grote, Fabrizio Mafessoni, Petra Korlević, Mateja Hajdinjak, Benjamin Vernot, Laurits Skov, Pinghsun Hsieh, Stéphane Peyrégne, et al. A high-coverage neandertal genome from vindija cave in croatia. *Science*, 358(6363):655–658, 2017.

SP Yip. Sequence variation at the human abo locus. *Annals of human genetics*, 66(1):1–27, 2002.
